# Inhibition of mTOR signaling by genetic removal of p70 S6 Kinase 1 leads to anxiety-like disorders

**DOI:** 10.1101/2020.03.25.007427

**Authors:** Muriel Koehl, Elodie Ladeveze, Caterina Catania, Daniela Cota, Djoher Nora Abrous

## Abstract

The mechanistic target or rapamycin (mTOR) is a ubiquitously expressed kinase that acts through two complexes, mTORC1 and mTORC2, to regulate protein homeostasis as well as long lasting forms of synaptic and behavioral plasticity. Alteration of the mTOR pathway is classically involved in neurodegenerative disorders, and it has been linked to dysregulation of cognitive functions and affective states. However, information concerning the specific involvement of the p70 S6 kinase 1 (S6K1), a downstream target of the mTORC1 pathway, in learning and memory processes and in the regulation of affective states remains scant. To fill this gap, we exposed adult male mice lacking S6K1 to a battery of behavioral tests aimed at measuring their learning and memory capabilities by evaluating reference memory and flexibility with the Morris water maze, and associative memory using the contextual fear conditioning task. We also studied their anxiety- and depression-like behaviors by respectively performing elevated plus maze, open field, light-dark emergence tests, and sucrose preference and forced swim tests. We found that deleting S6K1 leads to a robust anxious phenotype concomitant with associative learning deficits; these symptoms are associated with a reduction of adult neurogenesis and neuronal atrophy in the hippocampus. Collectively, these results provide grounds for the understanding of anxiety reports after treatments with mTOR inhibitors and will be critical for the development of novel compounds targeting anxiety.

## INTRODUCTION

The mechanistic (or mammalian) target of rapamycin (mTOR) is an evolutionary conserved serine/threonine protein kinase that plays a key role in regulating protein synthesis. The mTOR pathway integrates signals from nutrients, growth factors, and energy status to regulate many processes, including cell growth, proliferation, motility, and survival^1,2^. In neurons, the mTOR pathway modulates local translation of proteins at the synapse and therefore is critical for different forms of synaptic plasticity^3,4^. Taken together with its ubiquitous expression, it is thus not surprising that dysfunction of mTOR signaling represents a common hallmark in a wide variety of brain disorders, including autism, tuberous sclerosis, neurofibromatosis, fragile X or Rett syndrome, and neurodegenerative disorders, such as Parkinson’s, Alzheimer’s, or Huntington’s disease^5^.

mTOR therefore constitutes an attractive therapeutic target, and great effort has been made to determine its therapeutic indications. For instance, mTOR inhibitors such as rapamycin or its analogue everolimus are now approved for the treatment of various disorders including cancer, and as immunosuppressive drugs in solid organ transplantation. Furthermore, many preclinical and clinical studies are under way to test the efficiency and safety of mTOR inhibition in cystic diseases, neurodegenerative diseases or metabolic disorders^6^. However, the signaling pathways that are regulated by mTOR are complex and a considerable number of metabolic or physiological side effects have been described after treatments with inhibitors^6^. In particular, and consistent with the involvement of the mTOR pathway in synaptic plasticity and memory processing^7^, treatment with these inhibitors was found to affect cognition and affective states, with highly discrepant results.

On one hand, rapamycin treatment in humans and preclinical animal models may induce significant cognitive impairment^8,9^ and increase depressive-like and anxiety-like behavior^10–12^. The latter observation is consistent with mouse models of disorders that impact mTOR signaling in which abnormal anxiety-like behaviors have frequently been demonstrated^13,14^, and with anxious feelings reported by patients undergoing treatment with everolimus (https://www.webmd.com/drugs/2/drug-173864/everolimus-immunosuppressive-oral/details/list-sideeffects). On the other hand, everolimus treatment has been associated with significant improvements in memory and concentration functions and in mood and life quality as well as global psychiatric symptoms^15^. In agreement with this clinical dataset, rapamycin or everolimus treatment in adult mice was found to improve spatial learning and memory capabilities and decrease depressive- and anxiety-like behaviors^16,17^. Finally studies also reported that everolimus treatment did not affect learning and memory, and had no influence on depression- or anxiety-like behavior^18^.

Altogether, these data indicate that there is no clear correlation between activity of mTOR pathway and side effects such as cognitive deficits, anxiety or depression. Such discrepancy could be linked to the complexity and broadness of the mTOR network/signaling^19^. Indeed, mTOR acts in cells by forming two distinct complexes, respectively called mTOR complex 1 (mTORC1) and mTOR complex 2 (mTORC2). mTORC1 functions as a nutrient/energy/redox sensor; its effects are mediated by the phosphorylation of downstream proteins, such as the 70-kDa ribosomal protein S6 kinase 1 (S6K1), which in turn controls protein homeostasis. Whilst mTORC2 activates Akt/protein kinase B, which plays a central role in the control of cell metabolism, cell stress resistance and cytoskeleton regulation. As treatments with the mTOR inhibitors can affect one, the other, or both complexes^20^, different resulting effects can be expected. The development of new, more specific therapeutic tools to prevent these deleterious behavioral side effects thus depends on a better characterization of the involvement of the different effectors of the pathway in regulating cognition and affective states. We therefore investigated the consequences of blocking the mTORC1 pathway on cognitive and emotional behavior utilizing genetically-engineered mice deficient for S6K1^21^. Adult hippocampal neurogenesis was also assessed as a mechanistic substrate of the behavioral measures.

## MATERIAL AND METHODS

### Animals

Male S6K1-/- mice (henceforth named S6K1-KO) and their WT littermates were obtained and genotyped as described^22^. At eight weeks of age, animals were housed individually in standard plastic rodent cages and maintained on a 12h light/dark cycle (light on at 7am) with free access to water and food. Four batches of mice were used: Batch 1 (n=11 mice/genotype) was used for characterizing the impact of S6K1 deletion on memory abilities, anxiety-related behavior, exploratory behavior, and adult neurogenesis; Batch 2 (n=6 mice/genotype) was used to characterize depression-related behavior; Batch 3 (n=4 WT and n=6 S6K1-KO mice) was used to analyze the dendritic morphology of newborn dentate granule neurons; and Batch 4 (n=3 mice/genotype) was used to assess the dendritic morphology of all dentate granule neurons by Golgi staining. All experimental procedures have been carried out following the European directives of September 22, 2010 (2010/63/UE) and animal studies were approved by the ethical committee of Bordeaux (CEEA50; Dir 13105).

### General procedures

#### Measurement of memory abilities

##### Water maze

The apparatus was a white circular pool (150 cm in diameter) located in a room with various distal cues, and filled with water maintained at 20°C and made opaque by the addition of a non-toxic white cosmetic adjuvant. Data were collected using a video camera fixed to the ceiling of the room and connected to a computerized tracking system (Videotrack, Viewpoint) located in an adjacent room. The tracking system allowed the calculation of escape latency and path length.

###### Pre-training

mice (batch 1) received a three-step pre-training session. First, they were allowed to swim for 60 sec in the water maze without the platform. Then, they were placed upon the platform (16 cm diameter) raised at the surface of the water where they were required to stay for at least 15 sec. Finally, they were allowed to swim for a 30 sec period that was ended by a climbing trial onto the hidden platform (1.5 cm below water level). At the end of the pre-training, all mice swam actively and were able to climb onto the platform and stay on it for 15 sec.

###### Training with variable start positions

Mice were required to locate the hidden platform using distal extra-maze cues. They received 3 daily trials separated by a 5 minute inter-trial interval during which they were held in their home cages. A trial terminated when the animal climbed onto the platform or after a 60 second cut-off time. The starting point differed for each trial and different sequences of starting points were used day to day.

###### Training with constant start positions

Upon completion of the first training, platform location was changed to a different quadrant, and mice were required to find the hidden platform using constant starting points. Procedures were similar to the ones used for training with variable start positions. When performances reached a stable level, animals were tested to locate the hidden platform from a novel start position (1 trial).

##### Contextual fear conditioning

Conditioning took place in a transparent Plexiglas box (30 x 24 x 22 cm high) with a floor made of 60 stainless steel rods (2 mm diameter, spaced 5 mm apart) connected to a shock generator (Imetronic, Bordeaux, France). The box was cleaned with 70% ethanol before each trial. Animals (batch 1) were submitted daily for 3 days to a 5 min contextual conditioning session during which they freely explored the apparatus for 3 min upon which one electric footshock (0.7 mA, 50 Hz, 2 s) was delivered. Mice were then free to explore the cage for two more minutes. Freezing behavior was scored over the first three minutes preceding shock delivery by an experimenter blind to the genotype of mice.

To exclude a distorted nociceptive sensory perception of electric shocks, mice were submitted to a shock sensitivity protocol and tested in the hot plate test. The first test was carried out in the same conditioning chamber. Each mouse was administered seven 1s footshocks of increasing amplitude (from 0.1 to 0.7 mA) with an intertrial interval of 30 s. Two observers, blind to genotype, scored shock sensitivity based on three behavioral strategies: flinching, running/jumping and vocalizing. Scoring indicated the first shock intensity at which each reaction was detected. For the second test, which measures potential genotype-related differences in nociception, mice were placed in a Plexiglas box on the surface of a hot plate which was maintained successively at 49, 52, and 55°C. The stimuli were presented using ascending order of intensity at 30-min intervals. Latency for the mouse to raise and lick its paw or jump up was recorded. Mice were removed from the hot plate to prevent tissue damage if they did not respond within 30s.

#### Measurement of anxiety-related behaviors

##### The elevated plus maze (EPM)

was conducted in a transparent Plexiglas apparatus with two open (45 x 5 cm) and two enclosed (45 x 5 x 17cm) arms that extended from a common central squared platform (5 x 5 cm). The floor of the maze was covered with black makrolon and was elevated 116 cm above the floor. The test session began with the mouse individually placed on the center square facing an open arm. Animals (batch1) were allowed to freely explore the maze for 5 min (90 lux dim light). A camera connected to a computer was utilized to track the mouse path during the entire session (©VideoTrack, Viewpoint). Automatic path analysis measured time spent in and total number of entries into the open and closed arms. Standard measures of rodent anxiety were calculated: % time and % entry in the open arms compared to total time and total entries into any arm of the maze; in addition, total number of entries and total distance travelled in the open and closed arms were taken as a measure of activity/exploratory tendency in the EPM.

##### The open-field test

was used one day later as an additional measure of anxious-like behavior, as well as to evaluate locomotor performance and exploratory activity. It consisted of an illuminated square arena of 50 × 50 cm closed by a wall of 50 cm high and made in white PVC (light ~700 lux). Mice were placed individually in a corner of the arena and their activity was recorded for 10 min using a videotracking system (©VideoTrack, Viewpoint). Time spent and distance traveled in each zone (corners, periphery and centerfield) were recorded and analyzed.

##### The light/dark emergence test

was conducted in the same open-field containing a cylinder (10 cm deep, 6.5 cm in diameter, dark gray PVC) located length-wise along one wall, with the open end 10 cm from the corner. The day following open-field exposure, mice were placed into the cylinder and tested for 15 min under bright light conditions (1500 lux). Initial latency to emerge from the cylinder, defined as placement of all four paws into the open field, as well as total number of exits from the cylinder and total time spent inside the cylinder were analyzed.

#### Measurement of exploratory behavior

##### Locomotor activity

(batch 1) was recorded from 2 to 4 pm under dim light (50 lux) in racks of 8 activity cages (18.2 cm x 12 cm x 22 cm) made of transparent Plexiglas and isolated from the surrounding environment. Each cage was equipped with two beams of infra-red captors and infrared counts were computed via an electronic interface coupling each cage with an on-line computer (Imetronic, Bordeaux, France).

##### The novel object test

was conducted in the open-field described previously. Mice (batch 1) were allowed to freely explore the empty open-field for 30 min *(“habituation”* condition). After this phase, they were temporarily placed back into their home cage while an object (8cm in height and 7cm in diameter) was placed in the center of the open-field. Then animals were placed back into the open-field, now containing the cup *(“novel object”* phase), and tested for an additional 30 min. The time spent exploring the center of the open-field (target zone) in the presence and in the absence of the cup was measured.

#### Measurements of depression-related behaviors

In a different batch of animals (Batch 2), the influence of S6K1 deletion on depression-related behaviors was examined by measuring *avolition* (lack of motivation or inability to initiate goal-directed behavior) in the nest building and sucrose splash tests, *anhedonia* in the sucrose preference test, and *resignation/behavioral despair* in the Forced swim test (FST)^23,24^.

##### Nest building

A cotton nestlet was placed in each cage in the morning and nest quality was evaluated 24 hours later using the following criteria: Score 1: intact cotton square; Score 2: partially used cotton square; Score 3: scattered cotton; Score 4: cotton gathered in a flat nest; Score 5: cotton gathered into a “ball” with a small passage for entry of the animal.

##### Sucrose splash test

Ten days later, a high viscosity 10 % sucrose solution was sprayed on the coat of the mice to induce a self-grooming behavior. Latency to initiate the first grooming episode, as well as frequency and duration of grooming over a 5-min period was measured immediately after applying the solution.

##### Sucrose preference test

Two weeks later, mice were first habituated for 48h to the presence of two drinking bottles filled with tap water. They were then given, for 48h, a free choice between one bottle filled with a 4% sucrose solution, and the other with tap water. To prevent possible effects of side preference in drinking behavior, the position of the bottles was switched after 24h. The consumption of water and sucrose solution was estimated by weighing the bottles. Sucrose intake was calculated as the amount of consumed sucrose in mg per gram body weight, and sucrose preference was calculated according to the formula: sucrose preference = (sucrose intake)/(sucrose intake + water intake) × 100.

##### The Forced swim test (FST)

was performed ten days later by individually placing mice into a glass cylinder (height 25 cm; Ø 18 cm) filled with 26 °C water to a depth of 20 cm. Behavior was recorded for 6 min with a camera positioned to view the top of the cylinder. The latency to float and the duration of immobility were scored off-line by an experimenter unaware of the experimental groups. A mouse was judged to be immobile when it remained floating in an upright position, making only the movements necessary to keep its head above the water.

#### Thymidine analog injections

Animals from batch 1 were injected with Bromo-2’desoxyuridine (BrdU, 50 mg/kg dissolved in 0.9% NaCl, 1 daily injection during 5 days) one month after completion of the behavioral tasks.

#### GFP-retrovirus injections

GFP-encoding retrovirus was produced as previously described^25^. Mice from batch 3 were anesthetized with a mixture of ketamine (100mg/kg; Imalgene 1000, Merial) /xylazine (10 mg/kg; Rompun, Bayer HealthCare) and received 100 μl of a local anesthetic (Lidocaïne) under the skin covering the skull. They received a unilateral stereotaxic injection of the viral preparation (coordinates from Bregma: AP - 2, ML +/− 1.8, DV −2.2). Injections (1 microliter) were performed using a pulled microcapillary glass tube at a rate of 0.25 μl/min.

#### Immunohistochemistry and stereological analysis

One month after BrdU labeling (batch 1) or GFP injections (batch 3), animals were anesthetized and perfused transcardially with 0.1M phosphate buffered saline (PBS, pH 7.4), followed by 4% buffered paraformaldehyde (PFA). Brains were collected and post-fixed in PFA at 4°C for a week. Subsequently, 40μm-thick coronal sections were cut using a vibratome (Leica) and stored in cryoprotectant medium (30% ethylene glycol, 30% glycerol in KPBS) at −20°C before staining.

Free-floating sections were processed in a standard immunohistochemical procedure in order to visualize BrdU (1/1000, Accurate), doublecortin (DCX; 1:8000; Sigma), Ki67 (1:1000, Novocastra), or GFP (1/8000, Millipore) -labeled cells. Briefly, after washing in PBS, sections were treated with methanol and 0.5% H_2_O_2_ for 30 min. Sections were washed again in PBS before incubation with a blocking solution containing 3% normal serum and 0.3% Triton X100 in PBS for 45 min at room temperature. They were then incubated for 48h at 4 °C with the primary antibodies diluted in the blocking buffer. The following day, sections were incubated with biotin-labeled secondary antibodies diluted in PBS—0.3% Triton X100—1% normal serum, and immunoreactivities were visualized by the biotin–streptavidin technique (ABC kit; Dako) with 3,3’-diaminobenzidine (DAB) as chromogen.

The number of immunoreactive (IR) cells throughout the entire granule and subgranular layers of the left DG was estimated using the optical fractionator method^26,27^.

#### Golgi staining

A separate batch of animals (Batch 4) was perfused transcardially with 2% paraformaldehyde and 2.5% glutaraldehyde in 0.1 M PBS, pH 7.4. Coronal vibratome sections for Golgi impregnation (100μm) were treated with 1% osmium tetroxide in PB for 30 min. They were then placed in 3.5% potassium dichromate overnight, followed by 6 h in 2% silver nitrate solution. The sections were finally dehydrated in graded alcohols, infiltrated in epoxy resin, mounted, and coverslipped on glass slides^28^.

#### Morphometric analysis of GFP- and Golgi-labeled neurons

The overall dendritic tree of GFP-immunoreactive and Golgi dentate granule neurons was measured as previously described^28,29^. Briefly, the morphometric analysis was performed with a X100 objective using a semiautomatic neuron tracing system (Neurolucida; MicroBrightField, Colchester, VT, USA). Neurons were traced in their entirety, and area of cell body, number of dendritic nodes, and total dendritic length were calculated. To measure the extent of dendritic growth away from the soma and the branching of dendrites at different distances from the soma, a Sholl analysis^30^ was carried out.

#### Statistical analysis

All statistical analyses were performed with Statistica 12.0 software (Statsoft) and results are reported Table 1. Student t-tests were used for comparing genotypes in anxiety- and depression-related behavior as well as in adult neurogenesis; Two-way ANOVAs with genotype and session as main factors were used whenever repeated measures were recorded and followed by a Tukey post-hoc analysis when appropriate. Scores for pain threshold in response to brief ascending foot shocks and for nest building quality were compared by means of Mann–Whitney U-test. In each analysis, a value of p<0.05 was considered significant. All data are presented as mean + SEM.

**Table 1.**
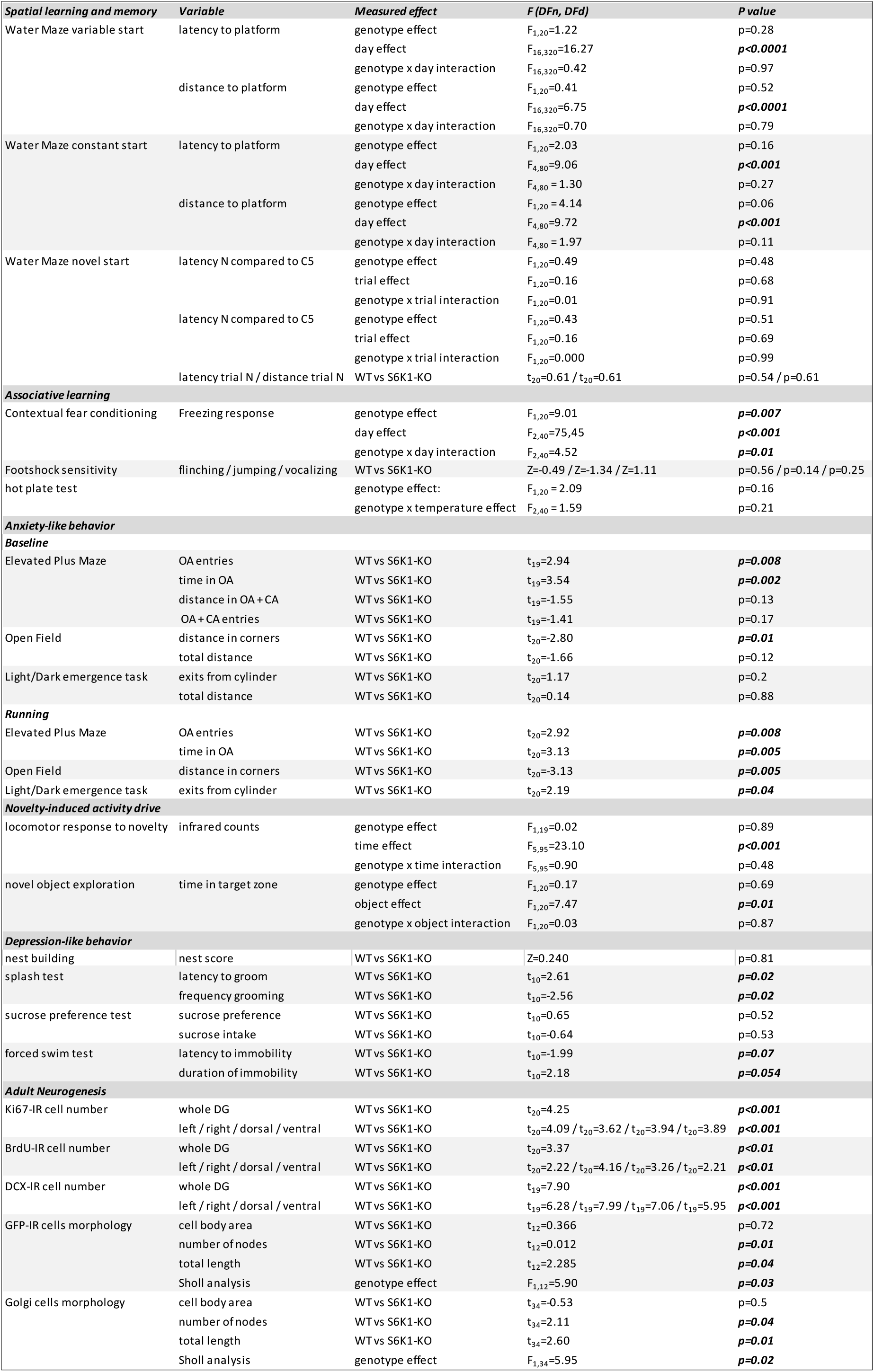
Statistics Table

## RESULTS

### Removal of S6K1 specifically alters contextual associative fear memory

We first examined whether removal of S6K1 impairs spatial learning and memory abilities by testing spatial navigation in the water maze. In this task, animals learn the location of a hidden platform using distal cues. It can be solved using multiple strategies in parallel, which requires the integrity of the hippocampus to different degrees. In the first procedure, the platform was maintained hidden (NW quadrant), and the starting point (NE, SW, or SE) was changed at each of the 3 daily trials. In order to find the hidden platform, the animal has to use an allocentric mapping strategy that consists of learning the positional relationships linking the cues (‘‘spatial relational memory’’). This relational representation is needed for using these cues in novel situations (i.e. changing starting position) and is consequently necessary to solve the task. This cognitive ability relies on the integrity of the hippocampus. Under these conditions, mice from both genotypes learned the platform position at a similar rate as seen by the diminution of latency and distance (Figure 1a, Table 1) necessary to find the platform. In the second procedure, the position of the hidden platform was changed (NW to NE) but the starting point was maintained constant for all trials (SW quadrant). In this case, although the development of a mapping strategy is not prevented, the animal can also learn the position of the platform using egocentric strategies consisting of, for example, the association of an invariant configuration of spatial cues to the escape platform (‘‘place learning’’). Egocentric strategies are very efficient for finding the platform if the starting point is maintained constant but fail to sustain the behavior if the starting point is suddenly changed. Under these conditions, mice of the 2 genotypes did not differ in the daily evolution of latency and distance to find the platform during the constant-start learning phase (Figure 1b C1 to C5, Table 1). When they were released from a new starting point at the end of the learning phase, all mice were able to find the platform, and performances did not differ between genotypes (Figure 1b trial N, Table 1). Taken together results of the 2 procedures confirm that mice of both genotypes present similar abilities in spatial memory and are able to develop an efficient relational strategy.

**Figure 1.**
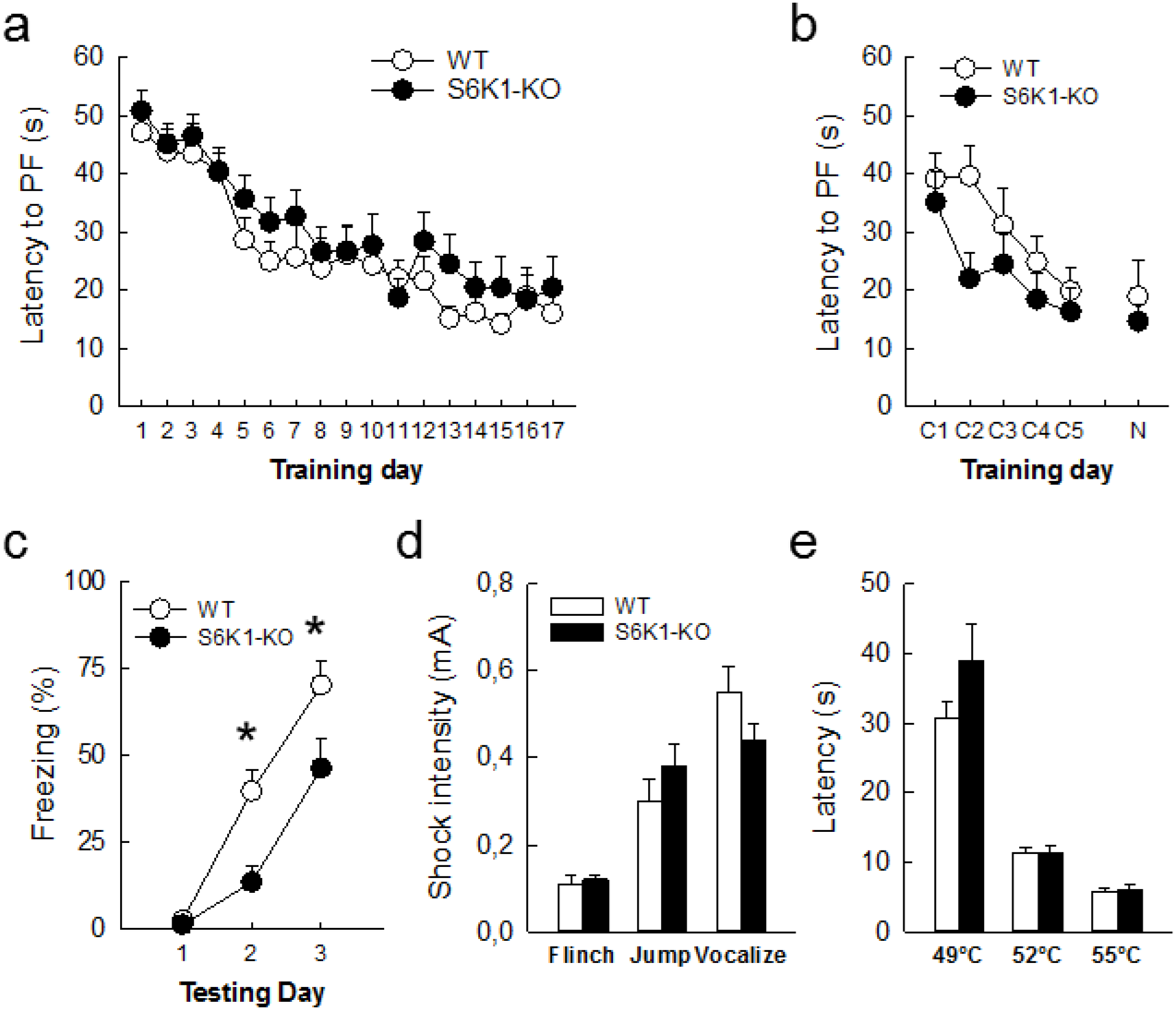
Removal of S6K1 spares spatial navigation but impairs contextual fear memory. **(a)** Latency to find the platform during reference memory testing **(b)** latency to find the platform location from constant start positions (C1 to C5) and a novel start position (N). **(c)** Freezing behavior (% total time) in response to a shock-associated context. (d) Threshold to elicit a flinch, jump or vocalize behavior in response to shocks of ascending intensity. (e) Latency to paw licking in response to ascending temperatures in the hot plate test. Data are mean ± SEM. n=11 mice per genotype. * p<0.05 compared to WT.

We then tested whether removal of S6K1 could alter the ability of mice to form and remember an association in the contextual fear conditioning task. If learning occurs, further exposure of an animal to the conditioning environment where it received an electric foot-shock elicits a freezing fear response. Mice (batch 1) received a single foot-shock each day during 3 days and their freezing response to the context-associated shock was recorded every day before shock exposure. Although both WT and S6K1-KO mice displayed an increased freezing across days, the latter reached much lower levels of freezing than WT (Figure 1c, Table 1). This difference was visible only on day 2 and day 3 (Tukey post-hoc test: day 1 p=0.99; day 2 p=0.01; day 3 p=0.03), indicating that the reduced freezing of S6K1-KO mice is not due to baseline differences but it is linked to a specific impairment in their ability to acquire a contextual associative fear memory. We further controlled that these differences were not due to an alteration of nociceptive sensory perception by measuring pain threshold in response to brief ascending foot-shocks or temperature setpoints. For both tests, the two groups did not differ (Figure 1d,e, Table 1), confirming that the decreased freezing observed in S6K1-KO mice is not linked to a lowered pain perception.

### Removal of S6K1 increases anxiety-like but not depression-like behavior

Anxiety-related behavior in rodents is mostly studied by measuring avoidance responses to potentially threatening situations, such as unfamiliar open environments. We first tested anxiety in the elevated plus maze (EPM) composed of two closed arms and two open arms, the latter constituting the threatening areas. Avoidance for these threatening areas was largely increased in S6K1 mutant mice, which visited less and spent less time in the open arms (Figure 2 a,b, Table 1). When exposed to a bright open-field (OF), again the behavior of the two groups was different as the distance travelled in the safest areas of the open field, the corners, was higher in mutant compared to control mice (Figure 2c, Table 1). Finally, mice were tested in the light/dark emergence task, a free exploration task in which animals can explore a brightly lit OF or retreat into a dark and reassuring cylinder. The number of exits from the cylinder, considered as an index of a lowered anxiety, was slightly decreased in the mutant mice, albeit this effect did not reach statistical significance (Figure 2d, Table 1).

**Figure 2.**
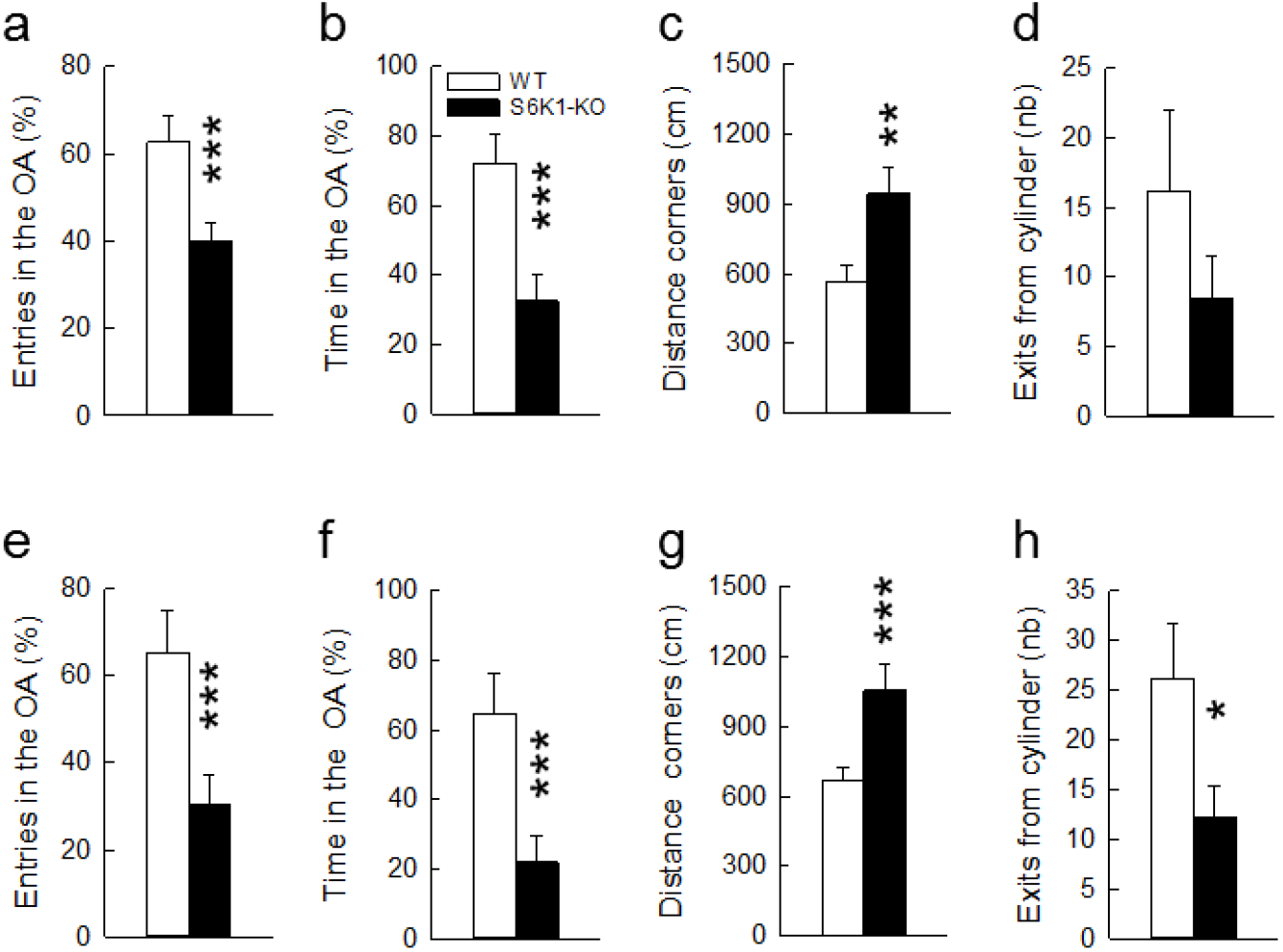
Removal of S6K1 increases anxiety-related behaviors. Anxiety-like responses were measured in the elevated plus maze (**a,b,e,f**), in the open-field (**c,g**) and in the light / dark emergence task (**d,h**) under baseline conditions (top) or after cage enrichment with a running wheel (bottom). Data are mean ± SEM, n=11 per genotype and per test except for the elevated plus maze under baseline conditions where one S6K1-KO mouse was removed as it fell from the maze. * p<0.05, ** p<0.01 and *** p<0.001 compared to the WT.

We verified that changes in activity/exploratory drive did not account for these phenotypic differences first by analyzing the exploratory tendency of mice in the different tests, then by measuring their activity drive in response to novelty in non-threatening situations. No differences in activity could be evidenced in the EPM (Distance travelled in both open and closed arms: WT: 7.95+0.9 m vs S6K1-KO: 10.31+1.2 m; Total number of entries in both open and closed arms: WT: 17.09+2.0, S6K1-KO: 21.5+2.3; Table 1), the OF (total distance travelled: WT: 21.9±1.5 m vs S6K1-KO: 31.8±5.7 m; Table 1), or the light/dark test (total distance travelled: WT: 23.9±3.6 m vs S6K1-KO: 22.9±5.5 m; Table 1). To test the activity drive in response to novelty, mice were tested for novelty-induced locomotor activity and novel object-induced exploratory activity in non-threatening environments. Both groups showed a similar decrease over time in locomotor activity as the context lost its novelty (Figure 3a, Table 1) and similarly explored the novel object, as shown by the increase in the time spent in the target zone when the object was present (Figure 3b, Table 1). These data indicate that the behavioral phenotype observed previously is not linked to an impairment in exploratory drive.

**Figure 3.**
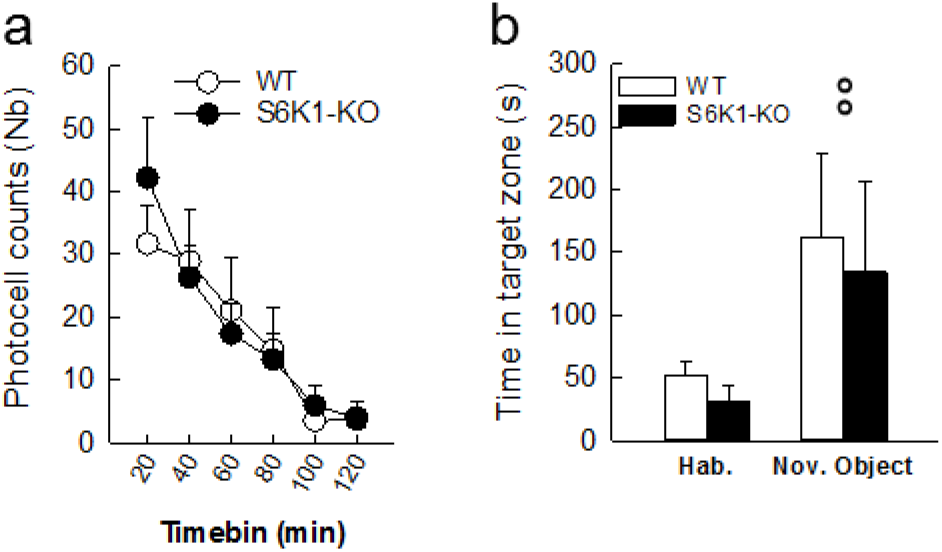
Removal of S6K1 does not alter activity drive in response to novelty. Locomotor activity was recorded in response to a novel environment **(a)** and exploration was recorded in presence of a novel object **(b)**. Data are mean ± SEM with n=11 per group, except for WT n=10 in response to novelty as one activity cage was deficient. °° p<0.01 compared to the habituation phase.

**Figure 4.**
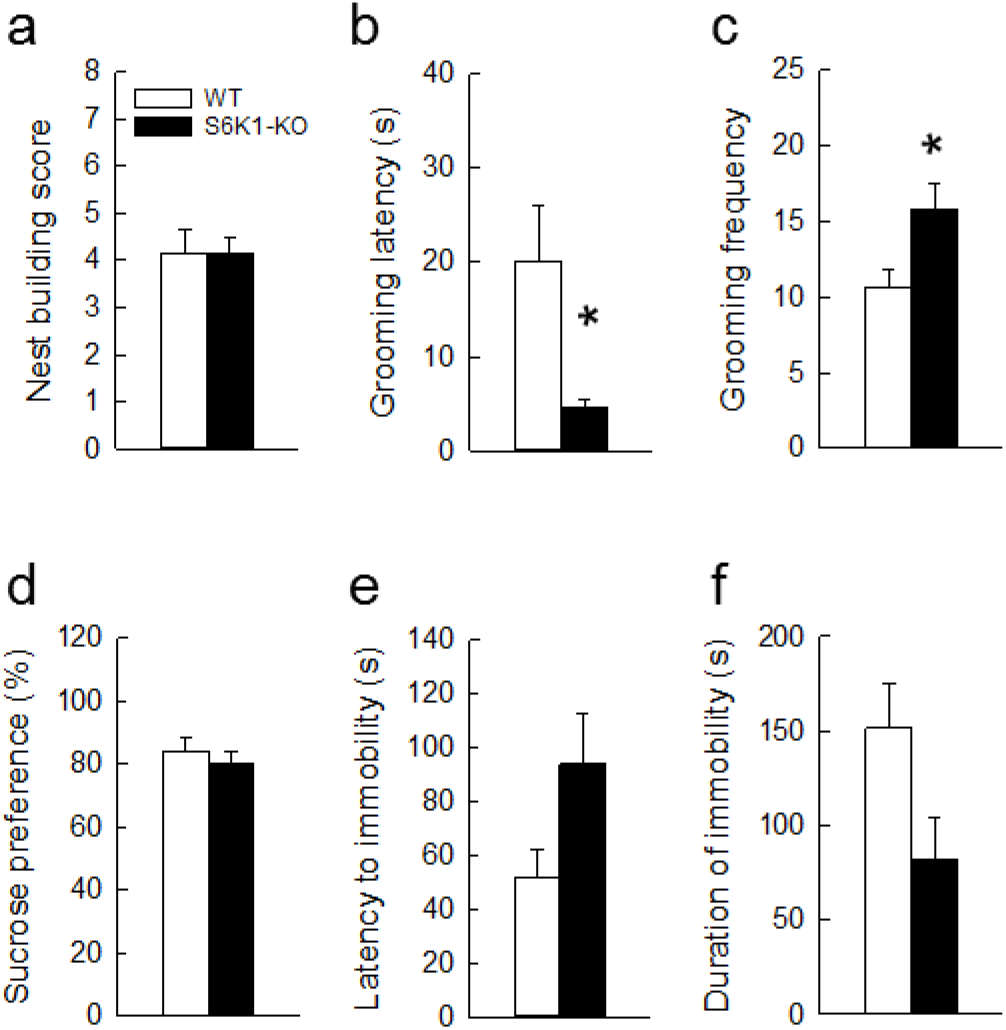
Removal of S6K1 does not increase depression-like responses. Motivation was evaluated in the nest building (a) and the sucrose splash test (b,c); anhedonia was measured in the sucrose preference test (d); and resignation was evaluated in the forced swim test (e,f). Data are mean ± SEM. n=6 per genotype. * p<0.05 compared to WT.

Then we asked whether anxiety-related behavior was stable and maintained when home cages were enriched by adding a running wheel, as running might bear anxiolytic potential^31^. Mice did not differ in their average daily use of the wheel (WT=1182+503 wheel revolutions; S6K1-KO=1142+470 wheel revolutions; t20=0.059, p=0.95). After 21 days of exposure to this new housing condition, anxiety-like responses were measured as previously done in the EPM (Figure 2e,f), the OF (Figure 2g) and the light/dark emergence task (Figure 2h). We found that the anxious phenotype was maintained and even more pronounced as differences between groups reached significance in the light/dark test (Table 1). Altogether these data indicate that anxiety-like behavior is an enduring feature of S6K1 mutant mice.

In the last series of experiments, the impact of removing S6K1 was examined on depression-related behavior using readouts of avolition *(nest building,* Figure 4a, *grooming in the splash test,* Figure 4b,c), anhedonia *(sucrose preference,* Figure 4d), or resignation *(forced swim test,* Figure 4e,f).

Nest quality was not different between groups (Figure 4a, Table 1) indicating that spontaneous motivation is spared following S6K1 deletion. Animals’ motivation toward self-centred activities was evaluated by measuring grooming behaviour. Typically, frequency and extent of grooming behavior is impaired in rodent models of depression leading to a degradation of coat states. Given that the coat states did not differ between the two groups of animals, we stimulated grooming behavior by splashing the back of the mice with a high viscosity sucrose solution. Opposite to what was expected, S6K1-KO mice developed an increased grooming behavior as shown by a decreased latency (Figure 4b, Table 1) and an increased frequency of grooming (Figure 4c, Table 1). In the sucrose preference test, in which a decrease in sucrose consumption is considered as an index of anhedonia, we found no differences between groups in both sucrose preference (Figure 4d, Table 1) and intake (WT: 11.4± 1.4 ml/g body weight; S6K1-KO: 12.9 ± 1.9 ml/g body weight, Table 1). In the forced swim test, although a strong tendency to increased active coping reflected by an increased latency to immobility (Figure 4e) and a deceased immobility time during the last 4 min of the test (Figure 4f) was recorded in S6K1-KO mice, this effect did not reach statistical significance (Table 1).

When analyzed together, this last dataset clearly indicates that S6K1-KO mice do not exhibit any consistent sign of anhedonia or “behavioral despair”. The analysis of each individual test wherein WT and S6K1-KO mice differ even suggests an increased elation-like behavior, but confounding factors cannot be excluded. Indeed, the excessive self-grooming displayed by S6K1-KO mice in the splash test could be linked to an increased propensity to obsessive-like response, but to the best of our knowledge, this type of behavior has never been tested in this model. As for differences in swimming behavior in the FST, buoyancy issues could be at play. Indeed, as previously reported^22^ S6K1-KO mice are smaller and we cannot exclude that decreased floating capabilities due to their lower body mass (WT m=43.55+1.4 g; S6K1-KO m=26.38+1.7 g; t10=7.52, p<0.001) translates into an increased swimming propensity in order to maintain flotation.

### Removal of S6K1 decreases adult hippocampal neurogenesis

Cell proliferation, examined using Ki67, was strongly decreased in S6K1-KO mice (Figure 5a, Table 1). This decrease was observed in both hemispheres and concerned both the dorsal and ventral parts of the dentate gyrus. This translated into a decreased number of 1-month old surviving BrdU-labeled cells (Figure 5b, Table 1) that was observed in both the left and right dentate gyrus, and concerned both the dorsal and ventral parts, as well as a decrease in the number of doublecortin-positive immature neurons (Figure 5c, Table 1) that was again observed in all sub-regions. Altogether, this indicates that removal of S6K1 decreases cell proliferation and alters adult neurogenesis. As the relevance of adult neurogenesis also relies on the synaptic integration of newborn cells, we checked the impact of S6K1 removal on the morphology of 4-wks old newborn neurons (Table 1). To this end, we used a GFP-encoding retrovirus that infects only dividing cells and allows cytoplasmic expression of GFP, thus providing a tool for dendritic analysis. Although the cell body area of newborn cells was not affected by the deletion of S6K1 (WT=98.28+6.9 μm^2^, S6K1-KO=94.78+6.4 μm^2^), all other parameters pointed to an atrophy of adult-born granule cells in S6K1-KO mice, which displayed less nodes (WT=6.16+0.5, S6K1-KO=2.75+0.9), shorter length (Figure 5d), and decreased complexity of dendrites (Figure 5d). Finally, in order to test whether this dendritic atrophy is restricted to adult-born cells or affects the entire population of granule neurons, we evaluated the morphology of neurons impregnated with Golgi on a separate set of animals (Table 1). As for newborn neurons, we found an atrophy (number of nodes WT=9.11+0.6, S6K1-KO=7.38+0.5; total length, Figure 5e) and a decreased complexity of dentate granule neurons in mutant mice (Figure 5e) without modifications of cell body area (WT=126.1±8.9 μm^2^, S6K1-KO=133.2±9.8 μm^2^).

**Figure 5.**
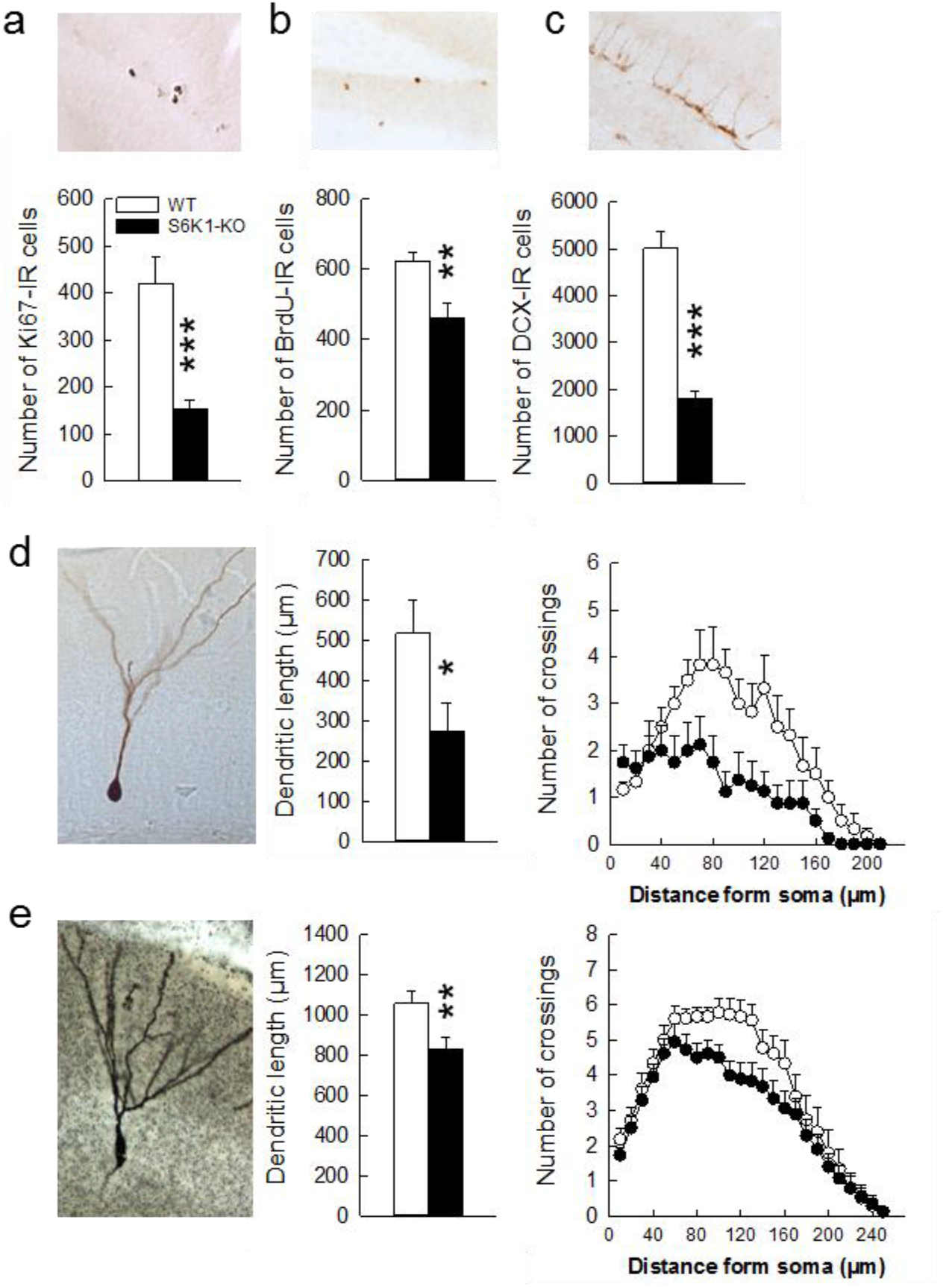
Removal of S6K1 decreases adult hippocampal neurogenesis and alters dentate granule cells morphology. **(a)** Cell proliferation, **(b)** Cell survival, **(c)** Neurogenesis, **(d)** Dendritic arborization of 4 weeks old adult born dentate neurons, **(e)** Dendritic arborization of dentate neurons impregnated with Golgi (n= 3 per group). Data are mean ± SEM with n=11 per genotype for a, b, c; n=6 WT and 8 S6K1-KO for d; n=18 neurons per genotype for e. *p<0.05, ** p<0.01 and *** p<0.001 compared to WT.

## DISCUSSION

There is a high comorbidity between neurodegenerative disorders, in which mTOR inhibitors are used as therapeutic approaches, cognitive defects and emotional dysregulation, such as anxiety and depression. Moreover, the mTOR pathway itself has been involved in learning and memory^7^ and recent studies have highlighted the involvement of mTORC1 signaling in stress-associated disorders, and in particular in depression^32^. The involvement of this pathway in anxiety has however remained elusive. Here we show for the first time that S6K1, a downstream target of mTORC1, mediates anxiety-like behaviors. Using a well-established transgenic mouse model, we confirm that S6K1 has a major impact on shock-induced contextual associative fear memory, and report unequivocally that removal of S6K1 increases anxiety-like behavior and seems to have anti-depressive actions. These alterations are associated with a defect in adult neurogenesis and a global atrophy of dentate neurons.

Studies that have so far investigated the role of mTORC1 signaling in regulating emotional states are sparse, and reports are controversial in regard to both anxiety- and depression-like behavioral effects. Comforting our own results on anxiety, it has been shown that exposure to a mild stress decreases hippocampal mTORC1 signaling and increases anxiety-like behavior^33^. On the same line, anxiolytic effects of fast-acting antidepressant drugs, such as YY-21, were reported to require activation of mTORC1 signaling in the medial prefrontal cortex (mPFC)^34^ and exercise, which can reduce the incidence of anxiety^35^, increases mTOR activity in the hippocampus and mPFC in rats^36^. Although this last dataset supports our findings, it was also reported that viral-mediated increased expression of S6K1 in the mPFC does not influence anxiety-like behaviors^37^. Furthermore, no differences in anxiety levels were found when testing the same genetic model of S6K1 deletion in an open-field^38^. In addition, pharmacological manipulation of the mTOR pathway with prescribed inhibitors also results in anxiogenic or anxiolytic effects, depending on the dose, route of administration, age of the subjects, and animal model used, as well as on possible preexisting neuropsychiatric predispositions or experimentally-induced neurological damage. For instance, chronic rapamycin treatment improved anxiety-like behaviors throughout lifespan in mice^17^, whereas treatment of male mice with the rapamycin analog everolimus induces anxiety-like behavior^11^. Consistent with this latter result, chronic treatment with rapamycin had anxiogenic effects in male rats^10,39^, as well as in a mouse model of Fragile X Syndrome, a neurodevelopmental disorder characterized by an upregulated mTORC1 signaling^40^. Finally, one study reported that rapamycin treatment in rats increased anxiety in a battery of tests without modifying phospho-S6K1 protein levels^10^. This observation led the authors to suggest that anxiety-related behavior after treatment with mTOR inhibitors could not directly be attributed to mTOR-dependent mechanisms. Our own data, directly testing the involvement of S6K1, contradict this hypothesis and strongly suggest that the anxiety induced by mTOR inhibitors can indeed be linked to an inhibition of the mTORC1 pathway. Altogether, the evidence currently available in the literature clearly point to a need to further investigate when and under which circumstances manipulation of mTORC1 signaling may differentially impact anxiety.

Although it was even less thoroughly tested, the involvement of mTORC1 in depression-like behavior is similarly controversial. We report that genetically blocking mTORC1 signaling does not induce depressive-like symptoms and even seems to have anti-depressive effects. This finding contrasts with previous studies indicating that reducing mTORC1 activity through 3-weeks treatment with everolimus^11^ or virally-mediated suppression of S6K1 activity in the mPFC of adult mice^37^ increases depressive-like behaviors. Our results are instead more in line with the observation that subchronic / chronic rapamycin treatment in both mice and rats exerts antidepressive-like effects^16,17,41^. Interestingly, a recent study reported that depression- and anxiety-like behaviors exhibited by mice in a Parkinson Disease (PD) model are eliminated by rapamycin, but not by selective blockade of the mTORC1 downstream target, S6K1^42^. Keeping in mind that these results were obtained in a PD-animal model, they can partly explain discrepancies in the existing data, and strongly corroborate the fact that inhibition of S6K1 does not recapitulate rapamycin actions. They also highlight the importance of gathering additional data on the consequences of manipulating downstream molecular targets of mTORC 1 in order to isolate potential candidate for medicating psychiatric symptoms both in baseline conditions and for comorbidities in neurological diseases where mTOR malfunctioning is manifest^10^.

In line with previous reports highlighting a role for mTORC1 signaling in both associative and non-associative fear memories^43^, our data also show that constitutive deletion of S6K1 causes a deficit in shock-induced contextual associative fear memory, while spatial navigation is spared. This dataset is consistent with reports that S6K1 is required for acquisition and consolidation of normal contextual fear memory but not necessary for spatial navigation^38^. The inability to acquire a shock-induced contextual associative fear was not due to an alteration of nociceptive sensory perception. Thus, we propose that an enhanced emotional reactivity linked to an anxious phenotype be at the origin of the associative fear deficits. Supporting this view, rapamycin administration blocks predator stress-induced associative fear memory^43^ as well as shock-induced inhibitory avoidance^3^.

In our studies, the increased emotional reactivity of S6K1-KO mice was associated with a reduction of adult neurogenesis. Cell proliferation was reduced in the DG, and as a consequence, the number of surviving cells and the number of immature neurons expressing DCX were also decreased. The complexity of the dendritic arbors of both adult-born and developmentally-born granule neurons was also altered as revealed by the diminution of dendritic length and complexity. Here again controversial data have been collected regarding mTOR and adult neurogenesis but the overall majority of studies seem to reach a consensus indicating that inhibition of the mTOR pathway decreases progenitor pools and neurogenesis^44–46^, which is in line with our own results. Only one study reported increased neurogenesis after chronic administration of everolimus^11^, and another one reported no effect of everolimus treatment on cell proliferation^18^. It should be noted that in this last study no behavioral consequences were observed after treatment and it is possible that their regimen of administration was subthreshold. Our data are also comforted by work from Dwyer et al. who reported increased dendritic branching of cortical neurons after transfection with a constitutively active form of S6K1^37^. Although the impact of inhibiting the activity of S6K1 was not tested, this is consistent with our own data and indicates that the relationships between S6K1 and neuron morphology extend to different structures. Although we did not directly test this hypothesis, it is highly conceivable that this alteration in adult neurogenesis could mediate the increased anxiety observed in S6K1-KO, as we have previously shown that decreasing hippocampal neurogenesis increases anxiety-like behavior^47^. This would also be consistent with the absence of depressive-like symptoms in mice deficient for S6K1 as we have also reported that reducing neurogenesis does not affect depressive-like behavior^47^. Therefore, the necessary and sufficient involvement of adult neurogenesis in driving anxiety in S6K1-deficient mice will definitely be worth testing.

Taken together our data point towards an important role for mTORC1 signaling, and more specifically S6K1 activity, in the regulation of anxiety-related behavior with two major consequences: first, these findings open new research avenues to evaluate whether pharmacological manipulation favoring S6K1 activity may favorably impact anxiety disorders, and second, they highlight the necessity to develop more specific pharmacological approaches to compensate for the side effects of prescribed mTOR inhibitors.

## Acknowledgments

We thank Sara Kozma and George Thomas (IDIBELL, Barcelona) for the kind gift of S6K1-KO mice. We thank the animal facility and the genotyping platforms of the INSERM U1215 NeuroCentre Magendie, funded by INSERM and Labex Brain, for animal care and mouse genotyping. The help of Delphine Gonzales and Cédric Dupuy is acknowledged. This work was supported by Institut National de la Santé et de la Recherche Médicale, INSERM (to DC and DNA), Aquitaine Region (to DC and DNA), the French National Research Agency, ANR (ANR-2010-1414, to DC and DNA; labex BRAIN ANR-10-LABX-43, to DC and DNA), European Community’s Seventh Framework Programme (Marie Curie IRG n°224757 to DC), and the Fondation pour la Recherche Médicale, FRM (post-doctoral fellowship to CC).

The sponsors had no role in the study design, collection, analysis and interpretation of the data, in the writing of the report, and in the decision to submit the article for publication.

